# Diverse and rapidly evolving viral strategies modulate arthropod vector gene expression

**DOI:** 10.1101/2021.12.16.472853

**Authors:** Nurit Eliash, Miyuki Suenaga, Alexander S. Mikheyev

## Abstract

Vector-borne viral diseases threaten human and wildlife worldwide. Vectors are often viewed as a passive syringe injecting the virus, however to survive, replicate and spread, viruses must manipulate vector biology. While most vector-borne viral research focuses on vectors transmitting a single virus, in reality vectors often carry diverse viruses. Yet how viruses affect the vectors remains poorly understood. Here we focused on the varroa mite, an emergent parasite that vectors over 20 honey bee viruses, and has been responsible for colony collapses worldwide, as well as changes in global viral populations. Co-evolution of the varroa and the viral community makes it possible to investigate whether viruses affect vector gene expression, and whether these interactions affect viral epidemiology. Using a large set of available varroa transcriptomes we identified how abundances of individual viruses affect the vector’s transcriptional network. Perhaps surprisingly, we found no evidence of competition between viruses, but rather that some virus abundances are positively correlated. Furthermore, we found a strong correlation between the extent to which a virus interacts with the vector’s gene expression and co-occurrence with other viruses, suggesting that interactions with the vector affect epidemiology. We experimentally validated this observation by silencing candidate genes using RNAi and found that reduction in varroa gene expression was accompanied by a change in viral load. Combined, the meta-transcriptomic analysis and experimental results shed light on the mechanism by which viruses interact with each other and with their vector to shape the disease course.

## Introduction

Vector borne viral diseases are infections caused by viruses and transmitted by infected arthropods (vectors) such as mosquitoes, ticks and mites (Brault et al., 2018; de la Fuente et al., 2017). They drive species evolution in both natural and man-managed ecosystems, and impose a great threat to organisms across kingdoms. Most disease-causing plant viruses depend completely on vectors for spread and survival (Ng & Falk, 2006; Whitfield et al., 2015), but animals also suffer from vector borne viruses (Olival et al., 2017). For humans, these include Dengue, Chikungunya, Chagas disease, Japanese encephalitis, Zika, and yellow fever, leading to hundreds of thousands of deaths worldwide especially in developing countries (Weaver et al., 2018; World Health Organization, 2017). Moreover, ongoing globalization processes and climate change are expected to increase their outbreak frequency (Rocklöv & Dubrow, 2020; Sutherst, 2004). The most efficient and sustainable measure of coping with emerging vector-borne diseases is to control their vectors (A. L. Wilson et al., 2020).

Vectors are often viewed as merely a passive syringe injecting the virus. However, to promote replication and transmission viruses must regulate the vector’s immune system and behaviour, which requires interaction with the vector. Indeed, former studies showed that viral infection can alter the vector feeding behaviour, fecundity, longevity and survival (Jackson et al., 2012; Maciel-de-Freitas et al., 2011; Moncayo et al., 2000; Neelakanta et al., 2010). The molecular mechanism underlying these interactions was studied in both cell cultures (Cime-Castillo et al., 2015; Göertz et al., 2019; Schnettler et al., 2014), and in vivo experiments (Luplertlop et al., 2011; Zink et al., 2015), revealing vector immune-related genes whose expression is regulated following viral infection, and show that a crosstalk between vector genes and the virus is imminent for successful viral replication and transmission (Huang et al., 2019).

Most of the studies have investigated the interaction of a vector with a single virus (Huang et al., 2019), and a few considered up to two co-occurring viruses (Goenaga et al., 2015; Göertz et al., 2017). However, vectors can usually carry diverse viruses that may interact with each other (Batovska et al., 2019; Batson et al., 2021; Ciota, 2019; Vogels et al., 2019). Studies on hosts infected by several viruses, have shown that coinfection affects viral traits such as virulence and transmission, either by direct virus-virus interaction, or indirectly by competition or cooperation (Erez et al., 2017; Ferguson et al., 2003; Nickbakhsh et al., 2019). Therefore, multi-infection may have a profound effect on viral evolution, diversity and pathogenicity (Alcaide et al., 2020; Díaz-Muñoz, 2017).

In contrast to these studies, the effect of multiple infections on vectors has received far less attention, although is expected to have considerable effect on the disease epidemiology (Ciota, 2019; Vogels et al., 2019). Therefore, virus-virus interaction is one of the main challenges in virology today (Sanjuán et al., 2021). We can expect that similar to the case of a host co-infected by several viruses, multi-infection will also have an effect on the vector-virus interaction. Yet we cannot simply infer from host-virus studies to vector-viruses, as the two are undergoing different evolutionary processes: while host-pathogen interactions are antagonistic by definition, vector-virus interaction are less definite and can fall anywhere on the continuum between antagonistic and mutualistic (A. J. Wilson et al., 2017). Therefore, we don’t know how vector-borne viruses interact, and how this may affect their relationship with their vector.

*Varroa destructor* (hereafter “varroa”) is a parasitic mite which vectors honeybee pathogenic viruses, and is routinely co-infected by multiple viruses (McMenamin & Genersch, 2015). The introduction of varroa to the European honey bee (*Apis mellifera*) has dramatically changed the honeybee viral landscape, leading to worldwide colony collapses threatening global food security (Steinhauer et al., 2018). In varroa-free colonies the viral diversity is high, with somewhat low levels (Carreck et al., 2010). Normally, one-two years following varroa introduction to a naive colony, viruses diversity and titers are shifting greatly, giving rise to high levels of a few viral strains as the colony is dwindling to its final collapse (Martin et al., 2012). Despite these observations, not much attention was given to varroa-virus interaction, and how it may explain the varroa key-role as a vector of honeybee viral disease. In addition, varroa carries over 20 diverse viruses commonly co-occurring (Table 1), including bee-pathogenic viruses, of which some are highly associated with varroa, while others are of unknown pathogenicity to either varroa or bee (Yañez et al., 2020). Therefore, the varroa-virus system is a unique opportunity to investigate how different viruses interact with each other and with their vector.

**Table 1.**
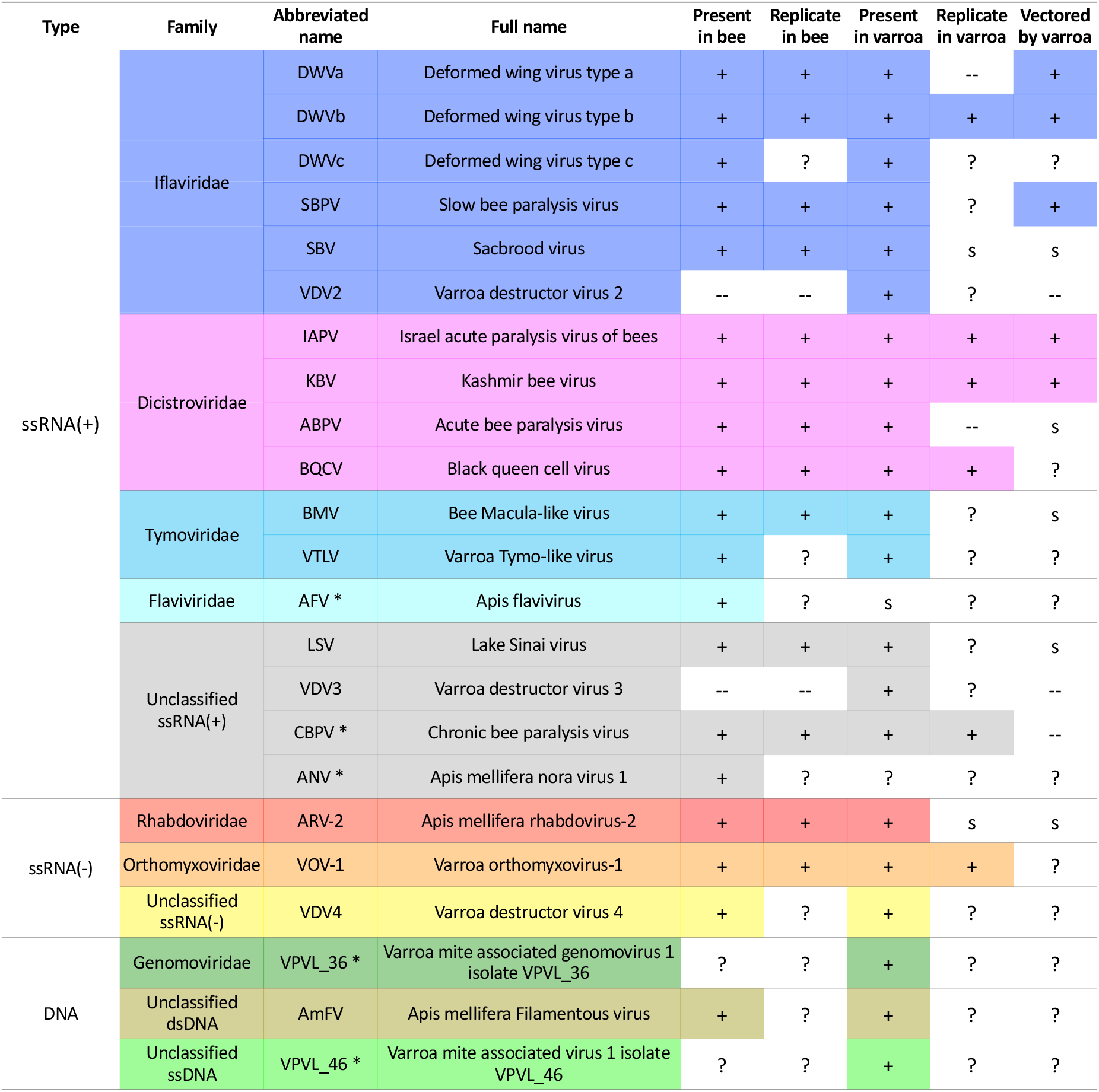
Most of the known bee viruses are present in both honey bee and mite, but only a few replicate in the mite, or shown to be varroa-vectored. Virus presence in bees and varroa, and its ability to replicate and vectored by the mite between bees are marked by: (+) confirmed; (--) not demonstrated; (S) suspected; (?) unknown. A star mark (*) next to the virus abbreviated name indicates that this virus was not detected in any of the varroa libraries in the current study. More details on each virus sequence and references can be found in Table S1.

Here we studied the interaction of diverse viruses with varroa’s transcriptional network, hypothesizing that different viruses have distinct effects. We explored a large set of RNAseq data from the vector aiming (1) to determine if viruses compete or cooperate in their vector, and (2) to test whether interactions between individual viruses and the vector correlate with viral epidemiological traits, specifically co occurrence with other viruses. We found no evidence of competition between viruses, but positive correlations in the abundance of specific viruses. Furthermore, we found a strong correlation between the extent to which a virus interacts with the vector’s gene expression and its overall prevalence, and have experimentally validated some of these predictions. These results show that multi-viral infection can occur not only in the host, but also in the vector, and that these interactions have implications on the virus relationship with its vector and therefore on the disease course.

## Results

### The viral landscape is heterogeneous across varroa libraries

We found that each of the final 66 varroa RNAseq libraries contain at least three types of viruses (Fig 1 and Table 1). The most prevalent viruses belong to the Iflaviridae family, including the bee pathogenic viruses DWVa and DWVb, and the varroa-specific virus, VDV2. On the other hand, five viruses whose presence in varroa was suspected or reported before, were not detected in any of the libraries (CBPV, AFV, ANV, VPVL_46 and VPVL_36) (Fig 1 and Table 1). Viral load homogeneity across libraries varied greatly between viruses: a few viruses were extremely variable (e.g, DWVa was not detected in a few libraries while reaching < 400,000 TPMs in others), while other viruses (such as ARV-2, VOV-1 and VDV2), were found in somewhat similar loads in most libraries. Interestingly, VDV2 is the only virus that was present in all varroa libraries.

**Figure 1.**
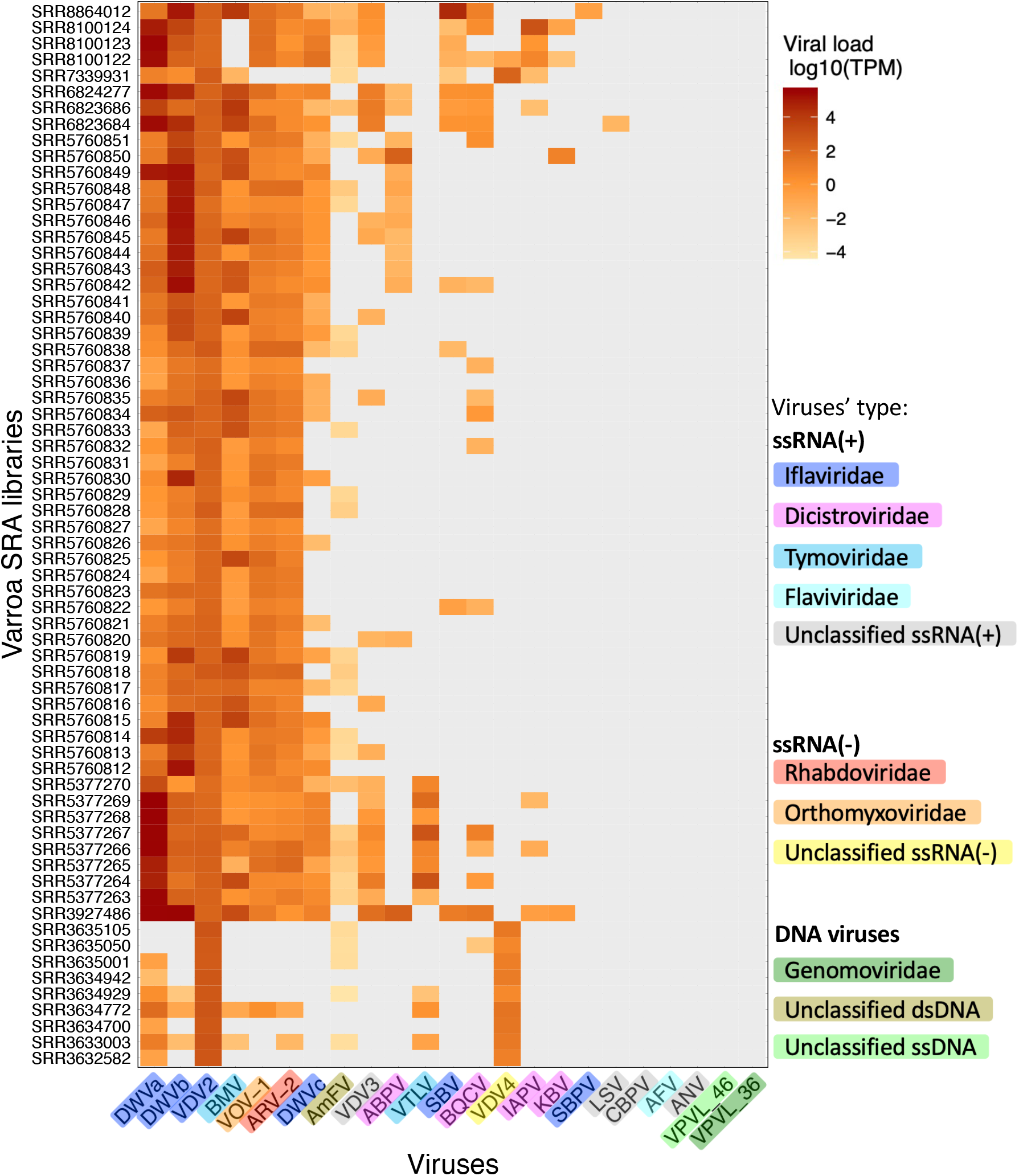
Viral load is diverse across the different varroa libraries. Members of the Iflavirus family are the most prevalent, yet while some viruses are homogenous across libraries (VDV2), others are highly diverse (e.g. DWVa and DWVb). Values are log10 transformed of the reads’ TPM (transcript per millions). Zero values are marked in grey (i.e., none of the reads in this library mapped to this virus). The viruses’ names are abbreviated as described in Table 1.

### No evidence for competition between the different viruses

Multi-infection by several viruses raises the obvious option of interactions between them. Interestingly, all significant correlations between the viruses’ viral loads were positive (Pearson correlation followed by Benjamini-Hochberg FDR-correction, P < 0.1) (Fig 2a).

**Figure 2.**
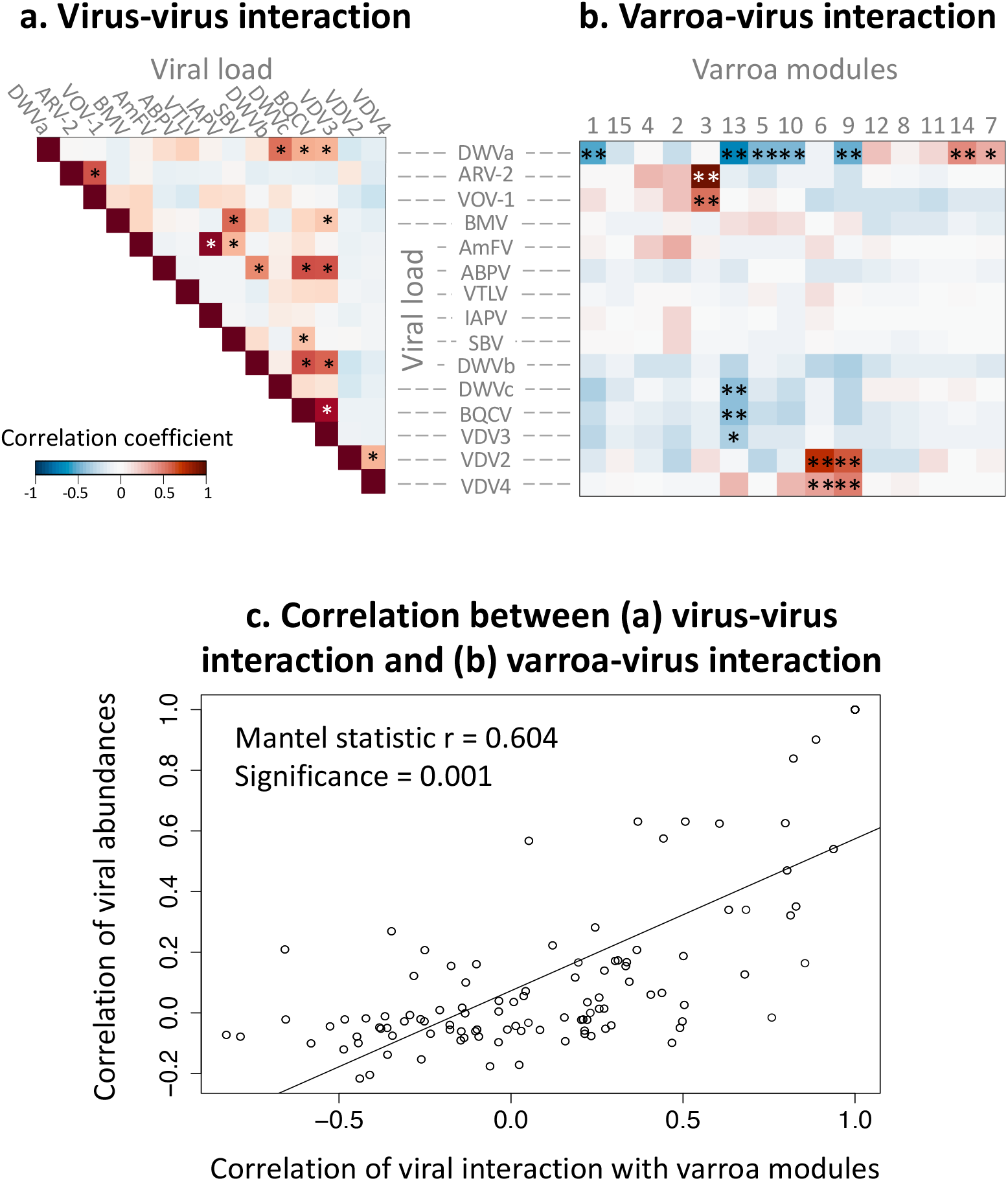
Intra-viral interactions can predict virus - vector interactions. a. Correlation between viruses’ loads. b. Correlation between viruses’ loads and varroa modules (eigengenes). c. Correlation model between viral load correlations (a), and the distance-matrix of the module-virus correlations (b). In figures a and b, viruses and modules are ordered according to hierarchical clustering; P values of the Pearson coefficient in figures a and b are adjusted according to FDR-correction; correlation significance marked by (*) 0.1 < P < 0.01; (**) P < 0.01. For analysis in figure c, Mantel test for correlation between two matrices was conducted using 1,000 permutations. All viruses’ loads are transformed by log10 of the (TPM), Transcripts Per Million.

### Varroa modules interact with its viral load

Gene network analysis of the 66 varroa RNAseq libraries clustered the 10,247 varroa genes into 15 modules, each module containing co-expressed genes. The modules were numbered from the largest (module number one which contains 3,625 co-expressed genes) to the smallest (module number 15 which contains 31 genes) (Fig S1a and b). Correlating varroa modules eigengenes to their viruses’ loads (Transcript Per Million, TPM), we found significant interactions between specific modules and viruses (Pearson correlation followed by Benjamini-Hochberg FDR-correction, P < 0.1, Fig 2b). The interaction direction and strength, represented by the correlation coefficient, varied between virus-module pairs. While VDV2, VDV4, ARV-2 and VOV-1 show positive interaction with some modules (modules 3, 6 and 9), ‘bee-viruses’ (such as DWVc and BQCV) show negative interaction with a different module (module 13). Interestingly, the only virus that shows both negative and positive interactions with vector’s modules is the bee-pathogenic virus, DWVa.

#### GO-terms of interacting modules

The significantly interacting modules consist of genes that are enriched for regulatory related GO-terms such as “Regulation of gene expression” (module 10), “Regulation of metabolic processes” (modules 6 and 9) and immune response related GO-terms, such as “Immunoglobulin production” GO:0002377 (module 6). In addition, modules 3 and 6 are enriched for viral and symbiont related GO-terms, such as “viral process” GO:0016032 and “modulation by virus of host process” GO:0019054, and “Regulation of biological process involved in symbiotic interaction”, GO:0043903. For the list of all significant GO-terms in the significantly interacting modules please see Tables S2-S10 (available on https://nurit-eliash.github.io/varroa-virus-networks/). Of the 17 RNAi-gene homologs recently identified in varroa (Beatrice T. Nganso et al., 2020), we found that 13 genes belong to modules interacting with viruses: modules 1 (negatively interacting with DWVa) and module 3 (positively interacting with ARV-2 and VOV-1) (Table S11).

### Viruses that are found together interact with the vector’s gene coexpression modules in similar ways

Among the module-virus interactions (Fig 2b), we can detect viruses that share a similar interaction with the same module. For example, VDV2 and VDV4 abundance positively correlated with modules 6 and 9. Interestingly, abundances of these two viruses also significantly correlate with each other (Fig 2a). Similarly, ARV-2 and VOV-1 also have positively correlated abundances and positive interactions with module 3 (Fig 2a and 2b). Likewise, DWVa, DWVc, BQCV and VDV3 are positively correlated in abundance, and all four are negatively correlated to module 13. We evaluated if this pattern between virus-virus interaction and virus-module interaction can be generalized across all virus-virus-module interactions in our data set. Indeed, we found a significant positive correlation between the two distance matrices (Mantel-test for correlation between two distance-matrices (Mantel, 1967), Mantel statistic r = 0.604, p = 0.001) (Fig 2c). In other words, there is a correlation between viral co-occurrence and the manner in which they interact with the vector.

### Validating varroa-virus interaction: gene silencing is accompanied by a change in viral load

To experimentally validate the varroa-virus interaction as predicted by the gene-network analysis, we silenced the mite’s genes using RNAi and tested its viral load, compared to the control (GFP-treated mites). For silencing, we selected candidate genes based on both the gene-network analysis and on the literature. From module 10, which interacts with the bee-pathogenic virus DWVa, we selected five which are highly connected to other genes in the module. In addition, these genes also possessed a relevant annotation, based on literature survey and / or the presence of conserved domain in the predicted coded protein (e.g. immune response related domains, and genes that were previously reported to interact with host/vector) (for the list of silenced genes, their module-membership and annotation, see table S12). Four of the five tested genes were successfully silenced, i.e., for these genes a significant reduction in relative gene expression was measured in mites treated with dsRNA, compared to control mites (P < 0.05, Wilcoxon signed-ranks test, followed by FDR-correction, table S13). We then compared the relative viral loads of DWVa, VDV2 and ARV-2 using quantitative PCR (qPCR). DWVa was predicted to negatively interact with the genes in module 10, while VDV2 and ARV-2 were not predicted to interact with the module (Fig 2b). Of all silenced genes, only mites that were treated with dsRNA of Cuticle protein type 8-like gene (short name: *CuP8*, accession: LOC111248360) showed a significant change in viral load, compared to control mites. Mites with decreased expression of *CuP8* had lower VDV2 and ARV-2 viral loads, compared to control mites (P = 0.02, Wilcoxon signed-ranks test, followed by FDR-correction), while DWVa viral load has also decreased, but not significantly (P = 0.2, Wilcoxon signed-ranks test, followed by FDR-correction) (Fig 3a and 3b).

**Figure 3.**
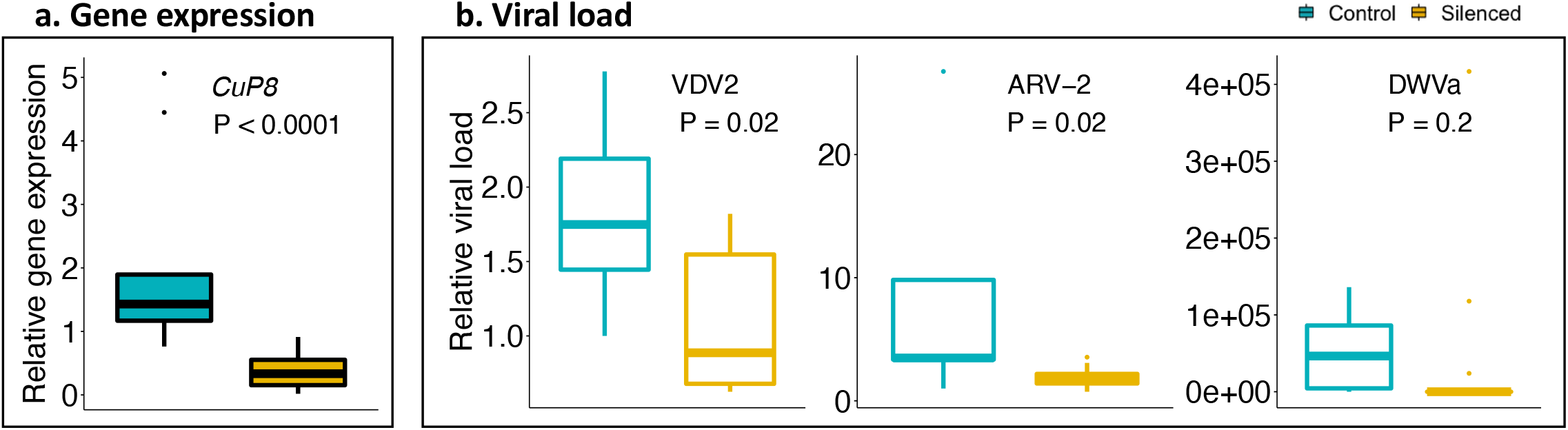
Validating varroa – virus interaction using gene-silencing (RNAi). **a**. Relative expression of *CuP8* gene in control (GFP-dsRNA, n=9) and silenced mites (treated with *CuP8*-dsRNA, n=13). **b**. Relative viral load of VDV2, ARV-2 and DWVa in control and silenced mites. Significant difference between control and silenced mites was tested using Wilcoxon signed-ranks test, followed by FDR-correction. The boxplot shows the interquartile of the data values, the inner thicker line of the box represents the median value, and the dots are potential outliers.

The dsRNA soaking treatment did not affect mite survival, compared to the control GFP-dsRNA treated mites (Fisher exact test for goodness of fit, p < 0.05) (Table S14).

## Discussion

Vector-borne viruses rely on another organism for transmission. As the vector plays a key role in viral fitness, we hypothesized that different viruses have distinct effects on a vector, observable as changes in gene expression. This was indeed true for viruses associated with varroa mites. Namely, titers of individual viral species were associated with specific changes to the vector’s gene expression network (Fig 2b). Interestingly, co-occurring viruses affected the vector gene expression network in similar ways (Fig 2c), suggesting a link between viral epidemiological traits and its relationship with the vector. We further propose that vector-virus interactions are evolving rapidly, as we found that closely related viruses have distinct effects on the vector’s gene expression. Interestingly, we found no evidence of viral competition within the vector. On the contrary, abundances of some viruses are positively correlated in co-infected vectors, suggesting a potential for viral cooperation. These complex dynamics underscore the role of vector-virus interactions for viral fitness.

### Viruses interact with specific modules in the vector’s transcriptional network

As we hypothesized, viral presence affects vector gene expression. We found that specific modules in the vector’s gene expression network respond to changes in viral titers in a species-specific way (Fig 2b). This indicates that the host responds to viral presence. Some of these responses may represent antiviral defenses (Göertz et al., 2019; Huang et al., 2019; Schnettler et al., 2014; Zink et al., 2015) or general stress responses (Rosche et al., 2021) that limit damage to the vector, though it is also possible that viruses trigger additional responses in a way that benefits their spread.

In accordance with this work, we found that the interacting modules involve both specific antiviral and non-specific stress responses. The modules included genes within the RNAi pathway (see Table S11), the main arthropod antiviral response (Blair & Olson, 2015). These modules were enriched for infection-specific GO-terms such as immune response and viral replication. At the same time, the modules were also enriched for regular cell maintenance GO-terms such as cell metabolism, and gene expression regulation. It should be noted that most of the modules responding to viral infection are not obviously related either to stress or antiviral gene expression, and their function *vis a vis* vector or viral fitness is unclear.

Following the network analysis, we have empirically validated the module-virus interactions using gene-silencing experiments of genes with high module connectivity, these are ‘hub-genes’ (Langfelder & Horvath, 2008). While we found that experimentally altering host gene expression does affect viral titre, the interactions were not in the direction the network analysis predicted. Namely we found that the knockdown of *CuP8*, a cuticular protein that has been found to assist in plant viral transmission by binding to viruses (Deshoux et al., 2018), was accompanied by a significant reduction in viral load of VDV2 and ARV-2, and non-significant reduction in DWVa (Fig 3b). Yet, *CuP8* was a hub-gene in a module negatively correlated with DWVa (Fig 2b), suggesting that it’s expression should be negatively correlated with that of the virus. The experimental result verifies on one hand the importance of the network hub-genes in the vector-virus interaction, while on the other hand, it illustrates the high complexity of the molecular mechanism underlying the vector response, which may involve other factors such as gene-to-gene transcriptional regulation, and interaction with the environment other microorganisms (Ciota, 2019).

### A link between viral epidemiological traits and their relationship with the vector

Interestingly, viruses with similar effect on the vector’s transcriptional network had tended to co-occur (Fig 2c). This observation can be explained by two non-mutually exclusive explanations. On one hand, high titres of viral RNA and proteins can trigger defense or stress responses by the vector (Baxter et al., 2017; Rosche et al., 2021). In addition, viruses could trigger these responses as a way to manipulate the vector’s ability to spread (Hurd, 2003; Targett, 2006). While the former mechanism is borne out by our data (see discussion of stress and antiviral pathways above), we can make several predictions about possible manipulation of vectors by the viruses. First, we would expect a more viral species-specific response by the vector, rather than a generalized response caused by increased virus titre. Second, we predict that the effect of the virus on the vector will evolve rapidly, with closely related viruses showing different effects on the vector. We explore these predictions below.

#### Species-specificity of vector-virus interactions

Viruses indeed show different patterns of interaction with varroa that seem linked to their ecology. For example, some of the more pathogenic bee viruses (DWVa, DWVc and BQCV) interact with the same module in the host gene expression network (Module 13, Fig 2b). These viruses are known to be associated with varroa infestation (Benjeddou et al., 2001; Daughenbaugh et al., 2015; Kevill et al., 2017; McMenamin & Genersch, 2015), and have a positive correlation with each other (Fig 2a). A different set of modules is affected by viruses that show a high level of expression in varroa, though are detected only at low levels in honey bees (VDV2, ARV-2, and VOV-1 (Fig 2a)) (Chen et al., 2021; S. Levin et al., 2016a, 2017), suggesting that these viruses may be infecting the mites, rather than using them as a vector. Interestingly, DWVa, the major driver of bee population declines, showed strong interactions with the varroa gene expression network, but modules associated with stress and antiviral responses were largely unaffected, suggesting that this virus may avoid them (Fig 2b). This could be due to the fact that the virus does not replicate in varroa (Gisder & Genersch, 2020), though it does trigger a number of other gene expression responses. These data suggest that viruses interact with varroa in diverse ways and that varroa gene expression is not likely solely driven by their titre.

#### Vector-virus interactions modified by a minute change in viral sequence

Virus interaction with its host can drive viral evolution and speciation (Koskella & Brockhurst, 2014), as host shift is a main force in viral diversification (Geoghegan et al., 2017; Ricklefs et al., 2014), and can change the pathogen virulence (Williams & Kamel, 2018). Our findings show that this also can occur in vector-virus interactions, as we found that closely related viruses have distinctly different effects on the vector’s gene expression. The two most well known variants of the DWV swarm, DWVa and DWVb are associated with varroa mites and colony collapses (Martin & Brettell, 2019). Varroa infection drives the DWV swarm evolution and the transition between the two variants (Dalmon et al., 2017; Gisder et al., 2018). Despite high sequence similarity, DWVb is less associated with the mite’s infestation level in the colony, compared to DWVa (Norton et al., 2021), while on the other hand DWVb is the only strain with concrete evidence for replicating in the mite (Gisder & Genersch, 2020). We therefore expected that the two variants, although highly similar in sequence, will have different interaction with the mite’s gene regulatory networks. Indeed, we found that the two variants show contrasting interactions with the vector modules: while DWVa shows both positive and negative interactions with the mite’s modules, DWVb shows no significant interactions with varroa modules (Fig 2b), which is surprising since DWVb was shown to replicate in the mite (Gisder & Genersch, 2020) and should therefore interact with the vector genes. However, we should note that the analysis can be biased by viruses with high titers, such as DWVa, which extreme high viral loads in many samples may mask other virus-module and virus-virus interactions. Our findings suggest that a small modification in the virus’s sequence can lead to a great change in the virus interaction with its vector, and that the vector-virus interactions are continuously and rapidly changing, resulting in a diverse viral community. Experimental infections of varroa by individual viruses can help further refine how individual viruses interact with the vector.

### Co-infecting viruses do not compete in the vector

The ‘competitive exclusion principle’ states that when competing in the same niche, one species will always suppress the other (Domingo, 2016). This was demonstrated also to happen in vectors, at least in mosquito cell-lines co-infected with two viruses (Abrao & da Fonseca, 2016; Pepin et al., 2008). In the light of the vector being a limited nourishing source, we could have expected the multi-infecting viruses to compete with each other in the varroa mite. However, we found exactly the opposite, and the only significant intra-viral interactions are positive ones (Fig 2a). This result, along with the high viral diversity in the varroa mite, support the model by Leeks *et al* (Leeks et al., 2018), which suggests that beneficial multi-viral interactions help to maintain high viral diversity. Still, our findings do not exclude the ‘short-sighted evolution’ model of virulence, which argues that in diverse infections, faster growing (more virulent) strains are favoured because they compete for limited resources (B. R. Levin & Bull, 1994). As the virulence of these viruses to varroa mite were not tested so far, we cannot conclude at this point the evolutionary mechanism which led to the observed multi-viral dynamic in the mite.

The positive correlation between some of the viruses may even imply mutualistic interactions, a phenomenon observed before for different strains of West Nile virus co-infecting mosquito (Ciota et al., 2012). A few mechanisms for mutualistic virus-virus relationships were suggested before such as cross-immunity, in multi-viral infection of influenza and other respiratory viral diseases (Ferguson et al., 2003; Nickbakhsh et al., 2019), and structural protein complementation in measles virus (Shirogane et al., 2012, 2016), and in mutants of Dengue virus infecting mosquito cells (Aaskov et al., 2006). However, the latter is the only example for vector-borne viruses. As viral colonization is the bottleneck for transmission, cooperative interaction between viruses in the vector have direct implications for the viral community dynamics, as they can favour specific viral strains that are not necessarily more virulent in a single, or even double infected vector. This further emphasizes the need to study multi-viral infections and their molecular mechanism.

Our study contributes to the ongoing investigation of the way viruses interact with their vector, and how this affects the disease course, specifically multi-viral infection, a current major gap in vector-borne viral research (Ciota, 2019; Vogels et al., 2019). Our results imply a link between the virus epidemiological traits and its relationship with the vector. In addition, not only that co-occurring viruses interact with each other, but their abundance may predict the way these viruses will regulate their vector, and potentially, its ability to successfully infect a new host. However, experiments on these viruses’ biology and effect on the vector are needed to have the full ecological context of these findings. Hopefully, the gene network pipeline established here can be adopted to other vector-borne diseases, opening the way to study the vector and its associated pathogenic and mutualistic symbionts (Hegde et al., 2015; Rainey et al., 2014).

## Materials and methods

In this study we investigated virus-virus and vector-virus interactions by two approaches: (1) meta-transcriptomic and (2) gene-silencing experiments. In the meta-transcriptomic analysis, we looked at the overall viral landscape in the different libraries, detected vector modules (co-expressing genes using gene-network analysis), and correlated the viruses’ abundance to the vector modules. Last, we tested if the virus abundance matrix can predict the virus-vector interaction. In the second step of the study, we experimentally validated the vector-virus interaction on specific virus-gene combinations, selected based on the meta-transcriptomic analysis, by RNAi-silencing varroa hub-genes and measuring the viral load. All analyses were carried out in the R statistical environment (Team, 2013). All meta-transcriptomic analyses are available and reproducible directly from the online supplementary data: https://github.com/nurit-eliash/varroa-virus-networks.

### (a) Vector-virus interaction, meta-transcriptomic analysis

#### Sequence Read Archive (SRA) data collection

To obtain varroa RNAseq data, we searched for “varroa” term in the SRA databases (NCBI, January 2020) with the following filtering criteria: “RNA” (Source filter), “RNA-seq” (Library Strategy filter) using Illumina technology, and the terms “TRANSCRIPTOMIC” and “VIRAL RNA” (Library Source filter). In total, the filtration yielded 71 libraries of varroa transcriptomes from 11 different studies. The libraries vary significantly in mite species, library preparation method and total library size, number of mites per sample, collected geographical region, sex, physiological stage, and the host species and developmental stage from which the mite was collected (for libraries details please see table S15).

#### Reads mapping and transcripts quantification

The reads were mapped to both available varroa genome (Vdes_3.0, accession number: GCF_002443255 (Techer et al., 2019)), and to the genomes of 23 selected viruses (Table 1). The alignment and estimation of transcript and virus abundances in transcripts per million (TPM), was carried out using Kallisto (Bray et al., 2016) (version 0.46.1 with default options). Kallisto pseudo-aligned reads to 35, 659 identified varroa isoforms from 10, 247 genes.

##### Mapping virus reads

The viruses were selected following a survey in the literature and NCBI genome database for viruses related to honey bee and/or varroa mite. Among the hundreds of virus sequences, we included only viruses that were previously detected in varroa, in addition to honey bee-pathogenic viruses (Table 1). The final 23 viruses are mostly positive ssRNA viruses, three are negative ssRNA viruses (ARV-2 (Rhabdoviridae), VOV-1 (Orthomyxoviridae) and VDV4 (unclassified)), and another three are DNA viruses (one circular dsDNA filamentous, and two ssDNA, found only in varroa mite and not in honey bees (Kraberger et al., 2018)). Although recently many DNA viruses were found in bees (Kraberger et al., 2019), their abundance and importance to varroa/bee health is unknown. Therefore, these sequences were not included in the current study.

#### Filtering data set

Given the diversity of sources, we wanted to make sure that the input data were as homogeneous as possible. A PCA of the different libraries based on varroa gene expression showed that five libraries are obvious outliers (Fig S2a). These were excluded from further analysis: library SRR8867385 (Brettell et al., 2019); libraries SRR5109825 and SRR5109827 from (Remnant et al., 2017), were deep sequenced for small RNA; and library SRR3927496 from a study by (Levin et al., 2016), in which a specific virome extraction procedure was implemented prior to library preparation. Last, library SRR533974 by (Cornman et al., 2013), was made of a pull of 1,000 mites, which is exceptionally higher than the number of mites used in most of the studies (a pool of one - five mites). All of these exceptional sample preparation procedures may account for these libraries’ deviation from the majority of the data sets in the PCA plot. The remaining 66 libraries are distributed somewhat homogeneously in a subsequent PCA (Fig S2b), and their reads were used for further analyses.

#### Viral abundance analysis

Viral abundance for all 23 viruses revealed that although reported before in varroa, five viruses were not detected in any of the libraires (CBPV, AFV, ANV, VPVl_46 and VPVL_36) (Fig 1), and so were not included in further analyses. To check for virus-virus interactions within the mite, we conducted correlation matrix abundance using Pearson correlation method for all viruses’ pairs on the log10 transformed TPM (Fig 2a).

#### Weighted gene network co-expression analysis (WGCNA)

We used a network analysis approach to identify groups of genes that share a similar expression pattern across a large set of available varroa transcriptomic data (RNAseq). To construct the gene-network, a weighted gene co-expression analysis was carried out using the WGCNA package in R, following the authors’ tutorial (Langfelder & Horvath, 2008, 2016). The WGCNA included 4 main steps: (1) Network construction and module detection; (2) Correlating modules to external information, the varroa viral load; (3) Identifying important genes; and (4) GO-term enrichment analysis for varroa modules which interact with the viral load.

##### (1) Network construction and module detection

Based on the analysis of network topology, for the construction of the network we set our threshold for merging of modules to 0.25, minimum number of 30 genes per module, and the power β of 12. This power is the lowest for which the scale-free topology fit index curve flattens out upon reaching a high value, in this case, when Rsq reaches 0.886 (Fig S1a). We then performed hierarchical clustering of the genes based on topological overlap (sharing of network neighborhood) to identify groups of genes who coexpressed across libraries, these are the network modules (Fig S1b).

##### (2) Correlating modules to viral load

To test if the varroa modules interact with the different viruses it carries, we correlated the module eigengenes to the viruses’ load (log10TPM). We used Pearson correlation method and adjusted the p-values for multiple comparisons using the Benjamini–Hochberg method to control the false discovery rate (Benjamini & Hochberg, 1995) (Fig 2b).

##### (3) Identifying hub genes in network modules

Hub genes have high connectivities within a module (Module Membership) (Langfelder & Horvath, 2008), and their annotation (based on sequence similarity to homologous genes). The Module Membership is calculated by Pearson correlation of the module eigengene and the gene expression. Genes with high Module Membership and relevant annotation are likely to play a role in the vector-virus interaction and are good candidates for later experimental validation.

##### (4) GO-term enrichment analysis for varroa modules

GO-terms enrichment analyses for the genes in the significantly interacting modules, were conducted with R package GOstats using the hypergeometric test for association of categories and genes (Falcon & Gentleman, 2007). The test parameters for each species and each ontology (biological process - BP) using gene ID from NCBI were as follow: p-value cutoff < 0.05, not conditional and with detection of over-represented GO terms (testDirection = over). All analyses are available and can be reproduced directly from the online supplementary data on GitHub: https://github.com/nurit-eliash/varroa-virus-networks.

#### Correlating virus interaction and virus-varroa interactions

To test if we can predict the virus-varroa interaction given the virus abundance, we used Mantel-test for correlation between two distance-matrices (Mantel, 1967).

### (b) Vector-virus interaction, mites’ gene - silencing experiment

In the second step of the study, we experimentally validated specific gene-virus interactions, as predicted by the meta-transcriptomic model. To check if indeed the gene expression is correlated to the varroa viral load, we silenced hub genes with high Module Membership and relevant annotation (see section ‘(3) Identifying hub genes in network modules’ in the meta-transcriptomic method part). We then checked for the viral load in the silenced mites. For the list of the selected genes for silencing, their accession and annotation please see Table S12.

#### Mites and honey bee collection

Mites and bees (*A. mellifera liguistica*) were collected from the same colonies, at the apiary of Okinawa institute of science and technology (OIST). The hives were not treated against mites and were supplemented with sugar solution and 70% pollen cakes as necessary. Mites were collected from drone and worker pupa of different stages and were kept on bees until soaking, up to five hours from collection.

#### RNAi silencing of varroa genes

##### DsRNA preparation

For the dsRNA preparation, we first synthesized a T7-promotor attached dsDNA of each of the targeted genes by PCR amplification of cDNA prepared from a pool of 5-10 mites, with specific primers with T7-promotor attached to the 5’ end (see Table S16 for the primers information, and section ‘varroa genes primer design’ for details on primer design). For PCR amplification we used Phusion™ High-Fidelity DNA Polymerase (Thermo Fisher Scientific) in a PTC-200 Peltier Thermal Cycler, MJ Research (BioRad, Toronto, Ontario) with the following steps: an initial denaturation at 98 °C for 30 secs followed by 30 cycles of denaturation at 98 °C for 10 sec, annealing at 60 °C for 10 sec, an extension at 72 °C for 30 sec, and a final extension of 72 °C for additional 5 mins. We checked that the size of the amplicons matches the expected length, by running 3 *μ*l of the PCR product in 1% agarose gel (135V, 20 min), then verified the sequence by purifying the product and Sanger sequencing on ThermoFisher SeqStudio Genetic Analyzer, using the original reverse primer and BigDye® Direct Cycle Sequencing kit (Thermo Fisher Scientific), following manufacturer’s instructions. Prior to dsRNA synthesis, we purified the PCR products using MinElute PCR purification kit (QIAGEN), measured their concentration by Qubit™4 Fluorometer (Life Technologies) with dsDNA HS Assay Kit (Invitrogen), and checked the product size by 4200 Tapestation (Agilent, Tokyo, Japan).

Next, 1200ng of the purified dsDNA with T7-promotor attached was used as a template for the dsRNA synthesis, using MEGAscript™ T7 Transcription Kit (Thermo Fisher Scientific). We followed the manufacturer’s protocol, with slight adjustments. The mix in total volume of 100ul was incubated overnight at 37 °C, following the addition of TURBO DNase buffer to the reaction, and incubation for another 15 mins at 37 °C. We purified and concentrated the RNA mix using MEGAclear Transcription Clean-Up kit (Thermo Fisher Scientific), and finally measured the RNA concentration by Nanodrop, and checked the product size by 4200 Tapestation (Agilent, Tokyo, Japan). To make sure that the dsRNA effect is specific, we also prepared a negative control dsRNA of a non-target gene, green fluorescent protein (GFP), using pET6Xhn-GFPuv vector (Clontech, Takara) as a template.

##### Soaking experiment

For applying the dsRNA into the mite body, we followed a protocol first developed by (Campbell et al., 2010), and successfully done by us and others (Garbian et al., 2012; B. T. Nganso et al., 2021; Singh et al., 2016). We added three mites in a 0.5ml tube containing 20*μ*l dsRNA (2.5*μ*g/*μ*l) in 0.9% NaCl solution. The tubes were kept in 4 °C for 10-15 mins then we checked that all of the mites were soaked, and re-dipped them into the solution, if needed. The mites were kept in 4 °C overnight (∼16 hours), dried on a filter paper, then put on a bee pupa (all same age), three mites per bee in a gelatin capsule with perforated top (#1, 0.49ml, HF capsules, Matuya, Japan). Following former studies (Campbell, Budge, et al., 2016; Singh et al., 2016), showing optimal silencing effectiveness 48 hours post dsRNA treatment, the mites were incubated for 48 hours in a controlled, dark environment at 34.5 °C, 60-75% RH, and the pupa was replaced after 24 hours. After incubation, each moving-viable mite was separated in a 1.5-ml tube, snap-frozen in liquid nitrogen and kept in -80 °C until RNA extraction. The experiment was replicated in seven experimental batches, between October - November 2020, and each batch included control group mites soaked in GFP-dsRNA of the same concentration and kept under the same conditions as described above. We checked the effect of the dsRNA treatment on mite viability, by comparing the numbers of live and dead dsRNA-treated mites to that of the control mites in each experimental batch (Fisher exact test for goodness of fit, p < 0.05) (table S14).

#### RNA extraction and cDNA preparation

Each individual mite was processed following the protocol developed in our lab and described previously (Hasegawa et al., 2021). Briefly, each individual mite was crushed in a 1.5 ml tube dipped in liquid nitrogen, then RNA was extracted using slightly modified TRIzol manufacturer’s protocol, with 50% volume of reagents. Total RNA quality and quantity were evaluated using Nanodrop spectrophotometer. 300ng of purified RNA was used to synthesize a first-strand cDNA using SuperScript II (Invitrogen) and 1:2 ratio of random hexamer and oligo dT primers following the manufacturer’s protocol.

#### Measuring varroa gene expression and viral load

For both viruses and genes, sets of primers were designed with NCBI primer design tool (utilizing Primer3 and BLAST), with default parameters and product size set to 100 - 400bp. Primer’s sequence, product length size and gene IDs or viruses’ accession numbers are provided in Tables S16 and S17 for varroa genes and viruses respectively. The product size of all amplicons was checked by running in 1% agarose gel (135V, 20 min).

##### Varroa genes primer design

For each of the varroa selected genes (Table S12) we designed two sets of primers, using the gene mRNA sequence as a template. A first set for dsRNA preparation (as described in ‘DsRNA preparation’ section), and a second, not overlapping primers-set for gene quantification using qPCR.

##### Identification of local viruses and RdRp primer design

We targeted the conserved gene of RNA-dependent RNA polymerase (RdRp) commonly used for detection and measurement of different RNA-viruses in honey bees and varroa mites (de Miranda et al., 2013; Gisder et al., 2018). As the genome sequence of bee viruses is slightly different across geographical regions (Cornman et al., 2013; Gisder et al., 2018; Gisder & Genersch, 2021), we first wanted to obtain the specific sequence of the strain present in our local mites. For that, for each of the three viruses (VDV2, ARV-2 and DWVa), we amplified and sequenced a wide region of the RdRp gene (amplicon size ∼800bp) (see section ‘DsRNA preparation’ for PCR and sequencing details). To verify the sequence, we nBlasted the product to the nr database (NCBI) (Altschul et al., 1990). The reverse complementary sequence of viral RdRp was then used as a template to design the qPCR primer sets, as described above. For viruses’ amplicon sequences, nBlast results and primer sequence and position, see Data S1.

##### qPCR

To evaluate the relative gene expression and viral load we performed a qPCR using a StepOnePlus Real-Time PCR system (Applied Biosystems Japan, Tokyo, Japan) with TB Green® Premix Ex Taq™ II (Tli RNaseH Plus, Takara). The cycling conditions were as follows: 95 °C for 30 secs, followed by 40 cycles at 95 °C for 5 seconds, and 60 °C for 30 secs. Data normalization and quantitating was done using StepOnePlus Real-Time system software (Applied Biosystems, Japan), with automatic threshold set. Small subunit of the ribosomal RNA gene (18S rRNA) of varroa was used as a normalizing gene (Campbell et al., 2016), and a control mite (treated with GFP-dsRNA) from the same experimental batch, used as the normalizing sample. For all qPCR assays a no-template control was included (data not shown). To test for differences in gene expression and viral load of dsRNA treated mites and control mites, we used a-parametric Wilcoxon signed-ranks test. For all statistical tests, the p-values were adjusted for multiple comparisons using the Benjamini– Hochberg method to control the false discovery rate (Benjamini & Hochberg, 1995).

## Acknowledgments

We wish to thank Dr. Olesya Gusachenko (University of St Andrews, UK) for consulting regarding the virus identification and viruses’ primer design. We also wish to thank Dr. Maeva Techer for her help in the tedious task of mites’ collection for the silencing experiment, and for all Mikheyev unit members at OIST for stimulating discussion and support throughout.

