## Supplementary information for "Diverse and rapidly evolving viral strategies modulate arthropod vector gene expression"

**Table S1.** Viruses mapped to the varroa RNAseq libraries in the meta-transcriptomic analysis. * In the original publication this virus was referred to as 'Varroa destructor virus-1', VDV-1 (Ongus et al., 2004), but today more often called DWVb. ** The genome of virus VOV-1 was published in six segments (Levin et al., 2019), therefore we mapped the varroa RNAseq reads to all six segments and summed their TPMs to account for VOV-1 viral abundance in each library.

| Type | Family | Short name | Full name | Accession number (NCBI) | Genome length (bp) | Full genome reference |
|---|---|---|---|---|---|---|
| ssRNA(+) | Iflaviridae | DWVa | Deformed wing virus type a | NC_004830.2 | 10,140 | (Lanzi et al., 2006) |
| | | DWVb * | Deformed wing virus type b | NC_006494.1 | 10,112 | (Ongus et al., 2004) |
| | | DWVc | Deformed wing virus type c | ENA\|CEND01000001\|CEND01000001.1 | 10,169 | (Mordecai et al., 2016) |
| | | SBPV | Slow bee paralysis virus | NC_014137.1 | 9,505 | (de Miranda et al., 2010) |
| | | SBV | Sacbrood virus | NC_002066.1 | 8,832 | (Ghosh et al., 1999) |
| | | VDV2 | Varroa destructor virus 2 | NC_040601.1 | 9,552 | (Levin et al., 2016) |
| | Dicistroviridae | IAPV | Israel acute paralysis virus of bees | NC_009025.1 | 9,499 | (Maori et al., 2007) |
| | | KBV | Kashmir bee virus | NC_004807.1 | 9,524 | (de Miranda et al., 2004) |
| | | ABPV | Acute bee paralysis virus | NC_002548.1 | 9,491 | (Govan et al., 2000) |
| | | BQCV | Black queen cell virus | NC_003784.1 | 8,550 | (Leat et al., 2000) |
| | Tymoviridae | BMV | Bee Macula-like virus | NC_027631.1 | 6,258 | (de Miranda et al., 2015) |
| | | VTLV | Varroa Tymo-like virus | NC_027619.1 | 6,169 | [Citation error] |
| | Flaviviridae | AFV | Apis flavivirus | NC_035071.1 | 20,414 | (Remnant et al., 2017) |
| | Unclassified ssRNA(+) | LSV | Lake Sinai virus | NC_032433.1 | 5,991 | (Daughenbaugh et al., 2015) |
| | Unclassified ssRNA(+) | VDV3 | Varroa destructor virus 3 | KX578272.1 | 4,202 | (Levin et al., 2016) |
| | Unclassified ssRNA(+) | CBPV | Chronic bee paralysis virus | NC_010711.1 | 3,674 | (Ribière et al., 2010) |
| | Unclassified ssRNA(+) | ANV | Apis mellifera nora virus 1 | KY354240.1 | 10,091 | (Remnant et al., 2017) |
| ssRNA(-) | Rhabdoviridae | ARV-2 | Apis mellifera rhabdovirus-2 | KY354234.1 | 14,001 | (Remnant et al., 2017) |
| | Orthomyxoviridae | VOV-1 | Varroa orthomyxovirus-1 | MK032465.1 | 2,198 | (Levin et al., 2019)** |
| | | | | MK032466.1 | 1,899 | |
| | | | | MK032467.1 | 1,983 | |
| | | | | MK032468.1 | 1,708 | |
| | | | | MK032469.1 | 1,442 | |
| | | | | MK032470.1 | 951 | |
| | Unclassified ssRNA(-) | VDV4 | Varroa destructor virus 4 | MK032464.1 | 8,332 | (Levin et al., 2019) |
| DNA | Genomoviridae | VPVL_36 | Varroa mite associated genomovirus 1 isolate VPVL_36 | MG571087.1 | 2,194 | (Kraberger et al., 2018) |
| | Unclassified dsDNA | AmFV | Apis mellifera Filamentous virus | KR819915.1 | 496,396 | (Gauthier et al., 2015) |
| | Unclassified ssDNA | VPVL_46 | Varroa mite associated virus 1 isolate VPVL_46 | MG571088.1 | 1,811 | (Kraberger et al., 2018) |

**Figure S1.** Network construction using 10,247 genes of 66 SRA varroa libraries. a. picking soft threshold. b. Hierarchical clustering dendrogram using merge cut height of 25%, revealing 15 co-expressed genes modules. Each branch of the dendrogram represents a single gene, and the colored bar below denotes its corresponding module, as annotated in the legend to the right. The dendrogram height is the distance between genes.

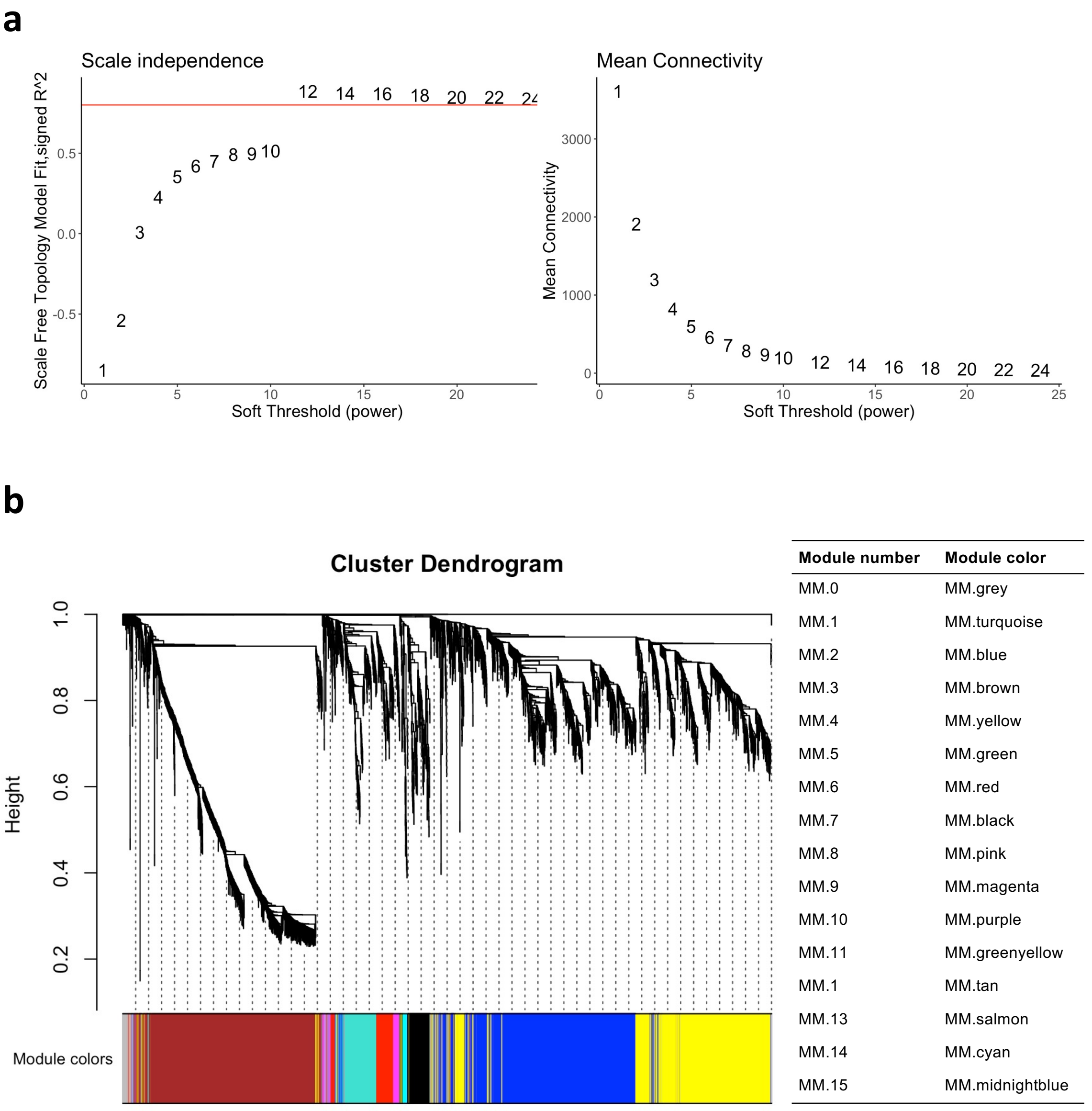

**Tables S2-S10** contain the significant GO-terms based on GO-term enrichment analysis (adjusted p-value<0.05), of nine modules. These modules were significantly interacting with at least one of the viruses (based on the analysis in Figure 2b). The tables are available on the online supplementary data: https://nurit-eliash.github.io/varroa-virus-networks/, in section "(4) GO-term enrichment analysis for varroa modules".

**Table S11.** Varroa RNAi pathway homolog genes, based on (Nganso et al., 2020), and the module it belongs to, based on the current network analysis.

| Family | Name | Protein | Gene | Module |
|---|---|---|---|---|
| **Dicer** | Vd-Dcr1 | XP_022665643.1 | 111252299 | 1 |
| **Dicer** | Vd-Dcr2a | XP_022645213.1 | 111243637 | 2 |
| **Dicer** | Vd-Dcr2b | XP_022645209.1 | 111243637 | 2 |
| **RdRp** | Vd1 | XP_022658093.1 | 111249053 | 1 |
| **RdRp** | Vd2 | XP_022666953.1 | 111252784 | 3 |
| **RdRp** | Vd3 | XP_022666954.1 | 111252784 | 3 |
| **RdRp** | Vd3 | XP_022647785.1 | 111244709 | 1 |
| **Argonaute** | Vd-Ago1 | XP_022655006.1 | 111247833 | 2 |
| **Argonaute** | Vd-Ago2a | XP_022665357.1 | 111252134 | 3 |
| **Argonaute** | Vd-Ago2b | XP_022656418.1 | 111248399 | 1 |
| **Argonaute** | Vd-Ago2c | XP_022671411.1 | 111254624 | 1 |
| **Argonaute** | Vd-Ago2d | XP_022650384.1 | 111245813 | 1 |
| **Argonaute** | Vd-Ago2e | XP_022646042.1 | 111243938 | 2 |
| **Argonaute** | Vd-Ago2f | XP_022672750.1 | 111255253 | 1 |
| **Argonaute** | Vd-Ago2g | XP_022656426.1 | 111248399 | 1 |
| **Argonaute** | Vd-Ago2h | XP_022656437.1 | 111248399 | 1 |
| **Argonaute** | Vd-Ago3 | XP_022669446.1 | 111253765 | 1 |

**Table S12.** Varroa hub-genes targeted for RNAi silencing.

| Gene ID | Gene description | Short name | Module Number | Module membership (Pearson correlation) | | Literature |
|---|---|---|---|---|---|---|
| | | | | Coefficient | P-adjust | |
| 111244103 | Glycerol-3-phosphate dehydrogenase | *Gly* | 10 | 0.86 | 3.4E-19 | Associated with a biotic stress and viral infection in both plants and humans (Kishimoto et al., 2012; Prasanth et al., 2011; Zhao et al., 2018). |
| 111244832 | Calmodulin | *clmd* | 10 | 0.85 | 1.4E-17 | An important factor for human cytomegalovirus replication (McArdle et al., 2011), and induce autophagy in rotavirus (Chattopadhyay et al., 2013; Crawford et al., 2012). |
| 111248360 | Cuticle-protein8 | *CuP8* | 10 | 0.77 | 5.8E-13 | cuticular proteins in the mouthparts of insect-vectors were found to bind plant-pathogenic viruses, thereby assisting in viral transmission while feeding on the plant-host (Deshoux et al., 2018). |
| 111245345 | Cuticle-protein-14 | *CuP14* | 10 | 0.61 | 7.0E-07 | |
| 111244631 | Twitchin-like | *Twitch* | 10 | 0.74 | 2.6E-11 | Contain consereved domains associated with immune-proteins (Immunoglobulin domain) |

**Table S13.** The probability that the difference between RNAi-treated and control mites (treated in GFP-dsRNA solution) have occurred by chance, for relative gene expression and viral load (Wilcoxon signed-ranks test, followed by FDR-correction).

| | | | | | Viral load | | |
|---|---|---|---|---|---|---|---|
| **Gene ID** | **Gene description** | **Short name** | **n** | **Gene expression** | **DWVa** | **VDV2** | **ARV-2** |
| 111244103 | Glycerol-3-phosphate dehydrogenase | Gly | 27 | 0.00 | 0.84 | 0.84 | 0.84 |
| 111244832 | Calmodulin | clmd | 24 | 0.00 | 0.84 | 0.84 | 0.84 |
| 111248360 | Cuticle-protein8 | CuP8 | 22 | 0.00 | 0.24 | 0.02 | 0.02 |
| 111245345 | Cuticle-protein-14 | CuP14 | 24 | 0.00 | 0.59 | 0.59 | 0.59 |
| 111244631 | Twitchin-like | Twitch | 17 | 0.09 | 0.68 | 0.68 | 0.68 |

**Table S14.** Fisher exact test of goodness of fit, testing the difference in live and dead mite distirbution between RNAi-treated and control mites (treated in GFP-dsRNA solution), in each experimental batch, ’Date’.

| Date | Silenced-gene | Live | Dead | pvalue |
|---|---|---|---|---|
| 08-Oct | Twitch | 9 | 0 | 1.00 |
| 08-Oct | Gly | 8 | 1 | 1.00 |
| 20-Oct | CuP14 | 4 | 5 | 1.00 |
| 21-Oct | Gly | 7 | 2 | 0.15 |
| 21-Oct | clmd | 2 | 7 | 1.00 |
| 23-Oct | CuP14 | 5 | 4 | 0.29 |
| 23-Oct | CuP8 | 8 | 1 | 1.00 |
| 23-Oct | clmd | 5 | 4 | 0.29 |
| 26-Oct | CuP8 | 5 | 4 | 0.33 |
| 17-Nov | clmd | 8 | 1 | 1.00 |
| 08-Oct | GFP | 9 | 0 | NA |
| 20-Oct | GFP | 4 | 5 | NA |
| 21-Oct | GFP | 3 | 6 | NA |
| 23-Oct | GFP | 8 | 1 | NA |
| 26-Oct | GFP | 2 | 7 | NA |
| 17-Nov | GFP | 8 | 1 | NA |
| 21-Dec | GFP | 12 | 6 | NA |

**Table S15.** Varroa SRA libraries used in the meta-transcriptomic analysis. The details in each column are based on the available information provided by the submitting authors on NCBI (https://www.ncbi.nlm.nih.gov/sra). When a detail is not provided in the database, we noted 'NS' for 'not stated'. Explanation for other short terms in the table: Am (*Apis mellifera*), Am Sy (*Apis mellifera syriaca*), Am In (*Apis mellifera intermissa*), Ac (*Apis cerana*), Am Cp (*Apis mellifera capensis*), Vd (*Varroa destructor*), Vj (*Varroa jacobsoni*).

| Library | Study | Mite stage | Bee species | Mite species | Collection method | Library selection | Other treatments |
|---|---|---|---|---|---|---|---|
| SRR6823684 | | Adult female | Am Sy | Vd | Adult bee | Random | mRNA enrichment |
| SRR6824277 | (Haddad et al., 2018) | Adult female | Am Sy | Vd | Adult bee | Random | mRNA enrichment |
| SRR6823686 | | Adult female | Am In | Vd | Adult bee | Random | mRNA enrichment |
| SRR5760851 | | Adult female | Am | Vd | Brood | cDNA | mRNA enrichment |
| SRR5760850 | | Adult female | Am | Vd | Adult bee | cDNA | mRNA enrichment |
| SRR5760849 | | Adult female | Am | Vd | Adult bee | cDNA | mRNA enrichment |
| SRR5760848 | | Adult male | Am | Vd | Brood | cDNA | mRNA enrichment |
| SRR5760847 | | Adult female | Am | Vd | Brood | cDNA | mRNA enrichment |
| SRR5760846 | | Adult female | Am | Vd | Brood | cDNA | mRNA enrichment |
| SRR5760845 | | Adult female | Am | Vd | Brood | cDNA | mRNA enrichment |
| SRR5760844 | | Adult female | Am | Vd | Brood | cDNA | mRNA enrichment |
| SRR5760843 | | Adult female | Am | Vd | Brood | cDNA | mRNA enrichment |
| SRR5760842 | | Adult female | Am | Vd | Brood | cDNA | mRNA enrichment |
| SRR5760841 | | Adult female | Am | Vd | Brood | cDNA | mRNA enrichment |
| SRR5760840 | | Adult female | Am | Vd | Adult bee | cDNA | mRNA enrichment |
| SRR5760839 | | Adult female | Am | Vd | Adult bee | cDNA | mRNA enrichment |
| SRR5760838 | | Adult male | Am | Vd | Brood | cDNA | mRNA enrichment |
| SRR5760837 | | Adult female | Am | Vd | Brood | cDNA | mRNA enrichment |
| SRR5760836 | | Adult female | Am | Vd | Brood | cDNA | mRNA enrichment |
| SRR5760835 | (Mondet et al., 2018) | Adult female | Am | Vd | Brood | cDNA | mRNA enrichment |
| SRR5760834 | | Adult female | Am | Vd | Brood | cDNA | mRNA enrichment |
| SRR5760833 | | Adult female | Am | Vd | Brood | cDNA | mRNA enrichment |
| SRR5760832 | | Adult female | Am | Vd | Brood | cDNA | mRNA enrichment |
| SRR5760831 | | Adult female | Am | Vd | Brood | cDNA | mRNA enrichment |
| SRR5760830 | | Adult female | Am | Vd | Adult bee | cDNA | mRNA enrichment |
| SRR5760829 | | Adult female | Am | Vd | Adult bee | cDNA | mRNA enrichment |
| SRR5760828 | | Adult male | Am | Vd | Brood | cDNA | mRNA enrichment |
| SRR5760827 | | Adult female | Am | Vd | Brood | cDNA | mRNA enrichment |
| SRR5760826 | | Adult female | Am | Vd | Brood | cDNA | mRNA enrichment |
| SRR5760825 | | Adult female | Am | Vd | Brood | cDNA | mRNA enrichment |
| SRR5760824 | | Adult female | Am | Vd | Brood | cDNA | mRNA enrichment |
| SRR5760823 | | Adult female | Am | Vd | Brood | cDNA | mRNA enrichment |
| SRR5760822 | | Adult female | Am | Vd | Brood | cDNA | mRNA enrichment |
| SRR5760821 | | Adult female | Am | Vd | Brood | cDNA | mRNA enrichment |
| SRR5760820 | | Adult female | Am | Vd | Adult bee | cDNA | mRNA enrichment |
| SRR5760819 | | Adult female | Am | Vd | Adult bee | cDNA | mRNA enrichment |
| SRR5760818 | | Adult male | Am | Vd | Brood | cDNA | mRNA enrichment |
| SRR5760817 | (Mondet et al., 2018) | Adult female | Am | Vd | Brood | cDNA | mRNA enrichment |

| | | | | | | | |
|---|---|---|---|---|---|---|---|
| SRR5760816 | | Adult female | Am | Vd | Brood | cDNA | mRNA enrichment |
| SRR5760815 | | Adult female | Am | Vd | Brood | cDNA | mRNA enrichment |
| SRR5760814 | | Adult female | Am | Vd | Brood | cDNA | mRNA enrichment |
| SRR5760813 | | Adult female | Am | Vd | Brood | cDNA | mRNA enrichment |
| SRR5760812 | | Adult female | Am | Vd | Brood | cDNA | mRNA enrichment |
| SRR3927486 | (Levin et al., 2016) | Adult female | Am | Vd | Adult bee | PolyA | mRNA enrichment |
| SRR3635105 | | Adult female | Am | Vj | Brood | cDNA | rRNA depletion |
| SRR3635050 | | Adult female | Am | Vj | Brood | cDNA | rRNA depletion |
| SRR3635001 | | Adult female | Am | Vj | Brood | cDNA | rRNA depletion |
| SRR3634942 | | Adult female | Am | Vj | Brood | cDNA | rRNA depletion |
| SRR3634929 | (Andino et al., 2016) | Adult female | Am | Vj | Brood | cDNA | rRNA depletion |
| SRR3634772 | | Adult female | Am | Vj | Brood | cDNA | rRNA depletion |
| SRR3634700 | | Adult female | Ac | Vj | Brood | cDNA | rRNA depletion |
| SRR3633003 | | Adult female | Ac | Vj | Brood | cDNA | rRNA depletion |
| SRR3632582 | | Adult female | Ac | Vj | Brood | cDNA | rRNA depletion |
| SRR8100122 | Shandong, China | Adult female | NS | Vd | NS | cDNA | mRNA enrichment |
| SRR8100123 | Shandong, China | Adult female | NS | Vd | NS | cDNA | mRNA enrichment |
| SRR8100124 | Shandong, China | Adult female | NS | Vd | NS | cDNA | mRNA enrichment |
| SRR7339931 | ARO, Israel | Adult female | Ac | Vd | NS | Random | NS |
| SRR5377270 | USDA-ARS | Egg | Am | Vd | Brood | Random | NS |
| SRR5377269 | USDA-ARS | Nymph female | Am | Vd | Brood | Random | NS |
| SRR5377268 | USDA-ARS | Adult female | Am | Vd | NS | Random | NS |
| SRR5377267 | USDA-ARS | Adult female | Am | Vd | NS | Random | NS |
| SRR5377266 | USDA-ARS | Nymph male | Am | Vd | Brood | Random | NS |
| SRR5377265 | USDA-ARS | Adult male | Am | Vd | Brood | Random | NS |
| SRR5377264 | USDA-ARS | Adult female | Am | Vd | Adult bee | Random | NS |
| SRR5377263 | USDA-ARS | Nymph female | Am | Vd | Brood | Random | NS |
| SRR8864012 | Valencia, Spain | Adult female | Am | Vd | Brood | PCR | NS |

*Outlier libraries (filtered out based on PCA (figure S1a)*

| **Library** | **Study** | **Mite stage** | **Bee species** | **Mite species** | **Collection method** | **Library selection** | **Other treatments** |
|---|---|---|---|---|---|---|---|
| SRR5109825 | (Remnant et al., 2017) | Adult female | Am Cp | Vd | NS | Random, Small RNA | rRNA depletion |
| SRR5109827 | | Adult female | Am Cp | Vd | NS | Random, Small RNA | rRNA depletion |
| SRR533974 | (Cornman et al., 2013) | Adult female | Am | Vd | Adult bee | Random PCR | NS |
| SRR3927496 | (Levin et al., 2016) | Adult female | Am | Vd | Adult bee | Random | virome |
| SRR8867385 | (Brettell et al., 2019) | Adult female | Am | Vd | Adult bee | cDNA | mRNA enrichment |

**Figure S2.** PCA of varroa SRA libraires based on their genes TPM. **a.** All initial 71 libraries. The outlier libraries are circled in red. These 5 libraries were excluded from further analysis. **b.** the final 66 libraries used for the analysis.

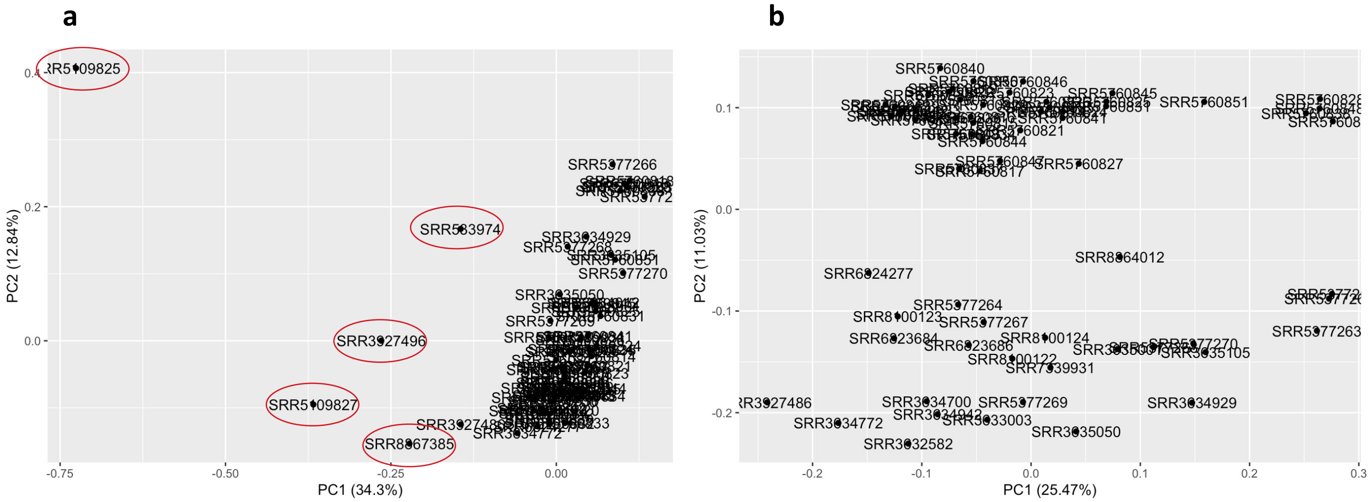

**Table S16.** Primers' sequences for varroa genes' primers for gene quantification using qPCR, and for dsRNA preperation. All primers are directed from the 5' to the 3' prime. For normalization of both varroa genes and viral abundacne, we used the small ribosomoal sub-unit, 18s as a reference gene.

| Gene description | Gene short name | Gene quantification (qPCR) | | |
|---|---|---|---|---|
| | | Primer sequence | | Product size (bp) |
| Small sub-unit of the ribosomal RNA (Campbell et al., 2016) | 18s | F | AATGCCATCATTACCATCCT | 60 |
| | | R | CAAAAACCAATCGGCAATCT | |

| Gene ID | Gene description | Gene short name | Gene quantification (qPCR) | | | dsRNA preparation | | |
|---|---|---|---|---|---|---|---|---|
| | | | Primer sequence | | Product size (bp) | Primer sequence | | Pr[illegible] |
| 111244103 | glycerol-3-phosphate dehydrogenase | Gly | F | CCAACTGTCGTTGCCCTATT | 128 | F | **TAATACGACTCACTATAGGG**CGAGGAGATCATCGATGGAAAG | [illegible] |
| | | | R | CACCGACGATTCTGGCTATTAC | | R | **TAATACGACTCACTATAGGG**CACTACGTCAGGGATAGCAATAA | |
| 111244832 | Calmodulin | clmd | F | CTGGGAAACCTGGCTGATAAA | 162 | F | **TAATACGACTCACTATAGGG**CGGAGCAAGAGCTCAAGAAA | [illegible] |
| | | | R | CCTTGCTCGTGCTAAGACTATC | | R | **TAATACGACTCACTATAGGG**CTACTTGGCGGAGGTGATAATG | |
| 111248360 | Cuticle-protein8 | CuP8 | F | CGACCTGAAACAAGCCATAGA | 256 | F | **TAATACGACTCACTATAGGG**GCCAAGGTGGATACGAATGA | [illegible] |
| | | | R | AGTGATACGGAGTCGGAGTAG | | R | **TAATACGACTCACTATAGGG**ACTTCATACGAGCGGGATTTAG | |
| 111245345 | Cuticle-protein-14 | CuP14 | F | TCAGTTAGTGCTTGCGTCTATG | 270 | F | **TAATACGACTCACTATAGGG**TCTACGCATTTCCGTCGTTATAG | [illegible] |
| | | | R | CAGCCAGCATAAGGGTGTATT | | R | **TAATACGACTCACTATAGGG**CGAGCGCGGTAAGTCAAATA | |
| 111244631 | Twitchin-like | Twitch | F | CGACACAGCACCGTGATAATA | 166 | F | **TAATACGACTCACTATAGGG**GGTTAGAGTTGGTGAGCCTATT | [illegible] |
| | | | R | GGAGAGTAGCCGACACAAATAC | | R | **TAATACGACTCACTATAGGG**CGATCCTGTCCACTGCTATTT | |
| | Non-target control gene green fluorescent protein | GFP | | | | F | **TAATACGACTCACTATAGGG**CGAAGTGGAGAGGGTGAAGGTGA | [illegible] |
| | | | | | | R | **TAATACGACTCACTATAGGG**CGAGGTAAAAGGACAGGGCCATC | |

**Table S17.** Primers' sequences for viruses RdRp sequencing and for viral load quantification using qPCR. All primers are directed from the 5' to the 3' prime. For normalization of viral abundacne, we used the small ribosomoal sub-unit, 18s as a reference gene.

| Accession number | Virus name | Virus short name | RdRp sequencing | | | Viral load quantification (qPCR) | | |
|---|---|---|---|---|---|---|---|---|
| | | | Primer sequence | | Product size (bp) | Primer sequence | | Product size (bp) |
| NC_004830.2 | Deformed wing virus, type a | DWVa | F | GCGTCCCGAACTTGAGATT | 893 | F | TCAACGACACAGTTAATGAGGA | 85 |
| | | | R | TCCAATTCGTCGTTCCTTCTAC | | R | TCCACAGGCAAACAAGTATCT | |
| NC_040601.1 | Varroa destructor virus 2 isolate VDV-2, complete genome | VDV2 | F | GGATCTGGAACATGCGATAGG | 759 | F | CAAGAGAATGGACAGACCTCTATG | 108 |
| | | | R | CGAGCACTCTCTTCAGACATTT | | R | CACCAATCTCAGTCGGAAGTT | |
| KY354234.1 | Apis mellifera rhabdovirus-2 (ARV-2) | ARV_2 | F | CCTAAGAGTGCAGTCCTTACAC | 893 | F | GGGAGTAGAAGGTTTGAGACAA | 150 |
| | | | R | GAGGTCCAGGTTTCGTCTATTT | | R | GGGTGTTTGTGGTACGGTAT | |

**Data S1.** Viruses' RdRp amplicon sequences, nBlast results and primer positions, for (a) DWVa, (b) VDV2 and (c) ARV-2.

### *(a) DWVa RdRp primer design*

Forwarded primer: 5' - GCGTCCCGAACTTGAGATT - 3'.
Reverse primer: 5' - TCCAATTCGTCGTTCCTTCTAC - 3'.
Amplicon (size 893bp) was used as a template to design a set of primers for viral quantification using qPCR. The qPCR amplicon product is underlined (85bp), and the qPCR primers' positions are highlighted, forward qPCR primer: 5' - TCAACGACACAGTTAATGAGGA - 3'; reverse qPCR primer: 5' - TCCACAGGCAAACAAGTATCT - 3'.

CCSRAACCTTGTMAGGGTTAGCCAGAAACACRGGTCTAGTTGGATGTTTTAAAAACCCGTGCTT
CAAGAAWGTAGCAGTCTGTAACGTCCGCCACTTCACAGTATTTCCTGATTTGTCCTGATCCGTA
AATTCCATCTTATATTGTGAAAAGAATTTCCCTATTGTCACAGCATTAAACTTATCRATCATGTT
GTCACTAACATTCATGATAAGATCATCACCATAACAAACAAGAACAACATTTTGAGAGAACTCG
GATAAAGGCAAATCAGTAATACCTAACCAAGCTAACCTAATTAACAGACAATTTGAAATYGTAT
TCAAAATGTCCGTTATCGGAGAACCTGATGGAATTCCACAAGGTACTCGGTACACTAAATCACGA
CATAGATGACTAGGCGCTAAAATCTCYTGCGCCATGGTCCACATTACTCGCTTCATTTCGTCTTT
ATTATCTTCTTCAGTRTAATGTAATACCCAGTCGATAATAATTTCRAACGCCGAAGCTGCAACAT
CGGAATCYAATCCAGGGCCAAAATTCTTATAGTCACCYGTCACGATATGAGTGCCATACTTTGAC
AAACTTGTTGCCAAATTTGTCCATTCTAAGCTGTTAACATCAATACCTATACCATGCTCAGCATT
AAGGCGTGCAGCTCGATAGGATGCCATAAAATCTAARTAATACTGTCTAAACGGTATAGTAAAC
TGTACTGGACTTATACTAAATATTCTAGTCTTACCAGGTATTCTACATTTTTCCACAGGCAAACA
AGTATCTTTCAAACAATCCGTGAATATAGTGTGAGGTTTATTCCCTTTTTCCTCATTAACTGTGT
CGTTGATACTGATCYAGAKTKYKGSRSMSRSMSC

nBlast first 10 hits, with lowest E value:

| Description | Scientific Name | Max Score | Total Score | Query Cover | E value | Per. ident | Acc. Len | Accession |
|---|---|---|---|---|---|---|---|---|
| Kakugo virus genomic RNA, complete genome | Kakugo virus | 1441 | 1441 | 96% | 0 | 97.04 | 10152 | AB070959.1 |
| Deformed wing virus isolate Varroa-infested-colony-DJE202, complete genome | Deformed wing virus | 1430 | 1430 | 96% | 0 | 96.8 | 10167 | KJ437447.1 |
| Synthetic construct clone DWVinfDVD5 polyprotein gene, complete cds | synthetic construct | 1419 | 1419 | 96% | 0 | 96.57 | 10264 | KT215904.1 |
| Deformed wing virus isolate VDV-1-DWV-No-5, complete genome | Deformed wing virus | 1419 | 1419 | 96% | 0 | 96.57 | 10149 | HM067437.1 |
| Deformed wing virus isolate Warwick-2009 polyprotein gene, complete cds | Deformed wing virus | 1419 | 1419 | 96% | 0 | 96.57 | 10167 | GU109335.1 |
| Deformed wing virus strain Liaoning-1, complete genome | Deformed wing virus | 1408 | 1408 | 96% | 0 | 96.33 | 10167 | MF770715.1 |
| Apis mellifera mRNA sequence | Apis mellifera | 1404 | 1404 | 96% | 0 | 96.32 | 1441 | HQ214486.1 |
| Kakugo virus gene for polyprotein, RNA dependent RNA polymerase region, partial cds, strain:MN3-050604 | Kakugo virus | 1393 | 1393 | 91% | 0 | 97.62 | 891 | AB242579.1 |
| Kakugo virus gene for polyprotein, RNA dependent RNA polymerase region, partial cds, strain:AM2-050526 | Kakugo virus | 1393 | 1393 | 91% | 0 | 97.62 | 891 | AB242578.1 |
| Deformed wing virus strain Korea-1, complete genome | Deformed wing virus | 1389 | 1389 | 97% | 0 | 95.76 | 10111 | JX878304.1 |

*(b) VDV2 RdRp primer design*

Forwarded primer: 5’ - GGATCTGGAACATGCGATAGG - 3’.

Reverse primer: 5’ - CGAGCACTCTCTTCAGACATTT - 3’.

Amplicon (size 759bp) was used as a template to design a set of primers for viral quantification using qPCR. The qPCR amplicon product is underlined (108bp), and the qPCR primers’ positions are highlighted, forward qPCR primer: 5’ - CAAGAGAATGGACAGACCTCTATG - 3’; reverse qPCR primer: 5’ - CACCAATCTCAGTCGGAAGTT - 3’.

AMRGRTTTGGMACTCTATTGGTTCATGCCCYTTYTTAATCCAATGTAGAATGCCAAACATTGAA
TCTTCAGCCAAGGGGGCAAGAAAATATACTCTTGAACTAATTAATTTTCTATCAGTAACCCTCTG
CCAATTACGTTTAAGRAAGGTCATATCATCTAATGTCCTATAYTTAACTGGAATCCCTTCCTTAT
CWGCATCAGTAAATTTTAAATCGTACCTGGCAAAGAAATCCCTAATTGTTAGCGTATTGAACAA
GTCACATACACAATCATCCAAGCCAATGATAACATCATCACCRTACGAGTACAAACTAGTATACY
TTTCAAAATTGTCCAACCCGCTAAATTCTGTGTTGGACATGATGGCCATCCATGCTATYCGAATA
AAAATGGAATTAACCATCGAATTTAAAACYACGGTCATGGTGTTACCACTTGGCAGTCCATTYAC
ACACTGRTACACCCAGCGGTCCGCAATGTGTTTAGCATTGAARACCTCTAGGGCCATAATAGAAA
GCACCCTTTGTATAATACACACCTGCTCYGGGGGTGCATACCGAGCATACCATGCACCAATCACA
CCAAACATCCGAACACCAATCTCAGTCGGAAGTTTGTCCCCAAATTTAGAATARTCCCCTGCTAT
AAATTTAGTTTTACCTCCTTTTGCTAATTTAACATAGAGGTCTGTCCATTCTCTTGAGTCCGGAT
TGATACCTATCGCWGTTYYYCMAAAWYMAMAA

nBlast first 10 hits, with lowest E value

| Description | Scientific Name | Max Score | Total Score | Query Cover | E value | Per. ident | Acc. Len | Accession |
|---|---|---|---|---|---|---|---|---|
| Varroa destructor virus 2 isolate VDV-2, complete genome | Varroa destructor virus 2 | 1042 | 1042 | 95% | 0 | 92.32 | 9552 | NC_040601.1 |
| Varroa destructor virus 2 strain NS, complete genome | Varroa destructor virus 2 | 1003 | 1003 | 95% | 0 | 91.33 | 9180 | MK795517.1 |

*(c) ARV-2 RdRp primer design*

Forwarded primer: 5’ - CCTAAGAGTGCAGTCCTTACAC - 3’.

Reverse primer: 5’ - GAGGTCCAGGTTTCGTCTATTT - 3’.

Amplicon (size 893bp) was used as a template to design a set of primers for viral quantification using qPCR. The qPCR amplicon product is underlined (150bp), and the qPCR primers’ positions are highlighted, forward qPCR primer: 5’ - GGGAGTAGAAGGTTTGAGACAA - 3’; reverse qPCR primer: 5’ - GGGTGTTTGTGGTACGGTAT - 3’.

CTGGCCMCCCGATAGATTTGAATTCTTCTTCAAGAGTGTCAAGAAAGAGTATTAACTTTCTTCG
GGCCTCATTCTCAGCAACTTCATTTGCCTCATCACAGTGAGGGTGTTTGTGGTACGGTATCCTCA
AACGAATTACCTGGTTATCACCCTGTCCAATGTTTTCAAAGGGAATTCGCAATTTTTCTGCAACT
TCCGTAATTTTACATACTGTTATAACAGTCCATGGCTTTTGTCTCAAACCTTCTACTCCCCCAAG
ATTACCTCTGAACCAGGAATCTCCTGGTATGAGATGACCATTCAAAAACTTGATAGTGTGTCCTT
CTCCTGTGTAAATCATTAGGCTTTTAGAAAAATGGTAATGTGTTTTTGACAAAACTCCACCRGG
ATAGCCCAGTATATCATCCATATCTTGGAAAAAAGGACGGACTAGAGGATCTCGGAAATTTGAG
TTCCATTTGGAGAAATCTATGTTGATTAATACATCAATACTTCTTTGGCGGATTCCCAAGGACCG
TGACATTCTGGCCATTAGGTTCCGAAGTTGATGTGCAGAATATGTCATAGTTACTTGTTTGAAG
TGAGGAAGTATATGTTCCTTAACTAGATGTTCTCCGATGACTGTCACCAATCTACAAGTGAGAGT
CTGTTGACCGAACATTCGAGCRGCAGGAAATTTCAGTTCCCTCTCCTTACAGGTTAGGATCATAA
TTTCATCTTCTGAGCTATACCCTTTCTCTTCTATCGAGCGTAAAAATTCCTTGGGGTTTGGGAAG
TTGGAGTCTAGAGCTTGCTCTATCAAACTCCTGCTATTTCTTGGTCCAGCATATCCCGTCTTTTC
AAGTTGTGTAGYMGCSCSCYCYYYYTMWWRAGRRA

nBlast first 10 hits, with lowest E value

| Description | Scientific Name | Max Score | Total Score | Query Cover | E value | Per. ident | Acc. Len | Accession |
|---|---|---|---|---|---|---|---|---|
| Apis rhabdovirus 2 isolate T-12 N protein, P protein, M protein, G protein, and L protein genes, complete cds | Apis rhabdovirus 2 | 1504 | 1504 | 97% | 0 | 98.47 | 14001 | KY354234.1 |
| Apis rhabdovirus 2 isolate RI-49 N protein, P protein, M protein, G protein, and L protein genes, complete cds | Apis rhabdovirus 2 | 1504 | 1504 | 97% | 0 | 98.47 | 14028 | KY354233.1 |
